# Self-assembly of hybrid 3D cultures by integrating living and synthetic cells

**DOI:** 10.1101/2024.10.05.616774

**Authors:** Nils Piernitzki, Ning Gao, Gilles Gasparoni, Julia Schulze-Hentrich, Michael Dustin, Balázs Győrffy, Stephen Mann, Oskar Staufer

**Author notes:** Corresponding author: Oskar Staufer.

## Abstract

Self-assembly, a fundamental property of living matter, drives the interconnected cellular organization of tissues. Synthetic cell models have been developed as bionic materials to mimic inherent cellular features such as self-assembly. Here, we leverage co-assembly of synthetic and natural cells to create hybrid living 3D cancer cultures. We screened synthetic cell models, including giant unilamellar vesicles, coacervates, microdroplet emulsions, proteinosomes, and colloidosomes, for their ability to form hybrid tumoroids. Our results identify the balance of inter- and extracellular adhesion and synthetic cell surface tension as key material properties driving successful co-assembly of hybrids. We further demonstrate that these synthetic cells can establish artificial tumor immune microenvironments (ART-TIMEs), mimicking immunogenic signals within tumoroids. Using the ART-TIME approach, we identify co-signaling mechanisms between PD-1 and CD2 as a driver in immune evasion of pancreatic ductal adenocarcinoma. Our findings demonstrate the 3D bottom-up self-assembly of hybrid cancer microenvironments to replace immune components with defined bionic materials, pushing the boundaries to functionally integrating living and non-living matter.

## Introduction

Cellular self-assembly into three dimensional architectures is a key process in life, critical for development and deregulated in cancer. Self-assembly of tissues is directed by intercellular signaling through biomolecular and biomechanical cues that result in spatially organized supra-cellular structures. Integrating this degree of self-organization and intercellular communication in man-made biomaterials, such as synthetic cells, is key to bridge the living and synthetic world. Self-assembling 3D cancer cultures, such as tumoroids and organoids, have been transformative in cancer biology research. These cultures, unlike their two-dimensional counterparts, emulate key biophysical and microanatomical aspects of natural tissues, such as extracellular matrix (ECM) networks, cavity formation, tissue asymmetry, spatial cell differentiation or metabolic gradients^1^. Specifically tumoroids, generated from cell lines or primary cancer cells, have facilitated substitution of animal models and unveiled novel disease mechanisms such as those involved in tumor immune evasion^2^. Tumoroid formation relies on the inherent self-assembly and self-organization properties of singularized cancer cells into 3D architectures. Several approaches have been developed to harness the intrinsic self-assembly capacity of cancer cells to form tumoroids, including matrix-assisted^3^, gravity-driven^4^, and restricted substrate adhesion procedures^5^. While the spatial organization in tumoroids is largely influenced by nutrient and growth factors gradients, their initial self-assembly into multicellular structures is an active process driven by *de novo* formation of intercellular adhesions during the aggregation process^6^.

Importantly, non-cancerous cells in the tumor microenvironment (TME) interact with cancer cells *in vivo*^7^, influencing tumor growth and therapy response, an interplay that is still challenging to replicate as self-assembling heterotypic tumoroids. The controlled and longtime integration of other cell types from the TME into tumoroids, such as fibroblasts, immune cells or neuronal cells, is limited as these do often not participate in the self-assembly process or are actively excluded and suppressed by cancer cells during the growth phase – a imitation that could be addressed by engineering biomimetic components.

In this study, we present an approach that stably integrates cellular bionic materials, in the form of synthetic cells into 3D cultures during self-assembly (**Fig. 1A**). This generates augmented tumoroids with artificial but controllable TMEs that bridge the natural and synthetic cell world. Synthetic cells have emerged as biomimetic materials that recapitulate molecularly defined characteristics of natural cells, allowing to study fundamental process of life such as cellular division, emergence of life and principles of cellular organization^8^. In this pursue, a whole toolbox of synthetic cell chassis has been developed, each designed to mimic individual biochemical and biophysical aspects of living cells. Giant unilamellar vesicles (GUVs) have been employed to replicate phenomena associated with cell membranes, such as pore formation and lipid phase separation^9^. Protein coacervates have been engineered to reconstitute protein-crowded environments within cells^10^ and inorganic colloidosomes have been developed to replicate the hierarchical organizational structures found within eukaryotic cells^11^. To specifically mimic cellular biomechanics, synthetic cells based on droplet-supported lipid bilayers (DSLBs), composed of a silicon oil core supporting a laterally mobile lipid bilayer (see also supplementary note 1), and polymerosomes have been designed^12–14^. Co-integrating synthetic cell models into self-assembling 3D natural cell constructs, could allow to systematically study the fundamental principles underlaying cellular self-assembly and open new avenues to apply synthetic cell materials as functional and controllable cell surrogates in 3D cell cultures.

**Figure 1.**
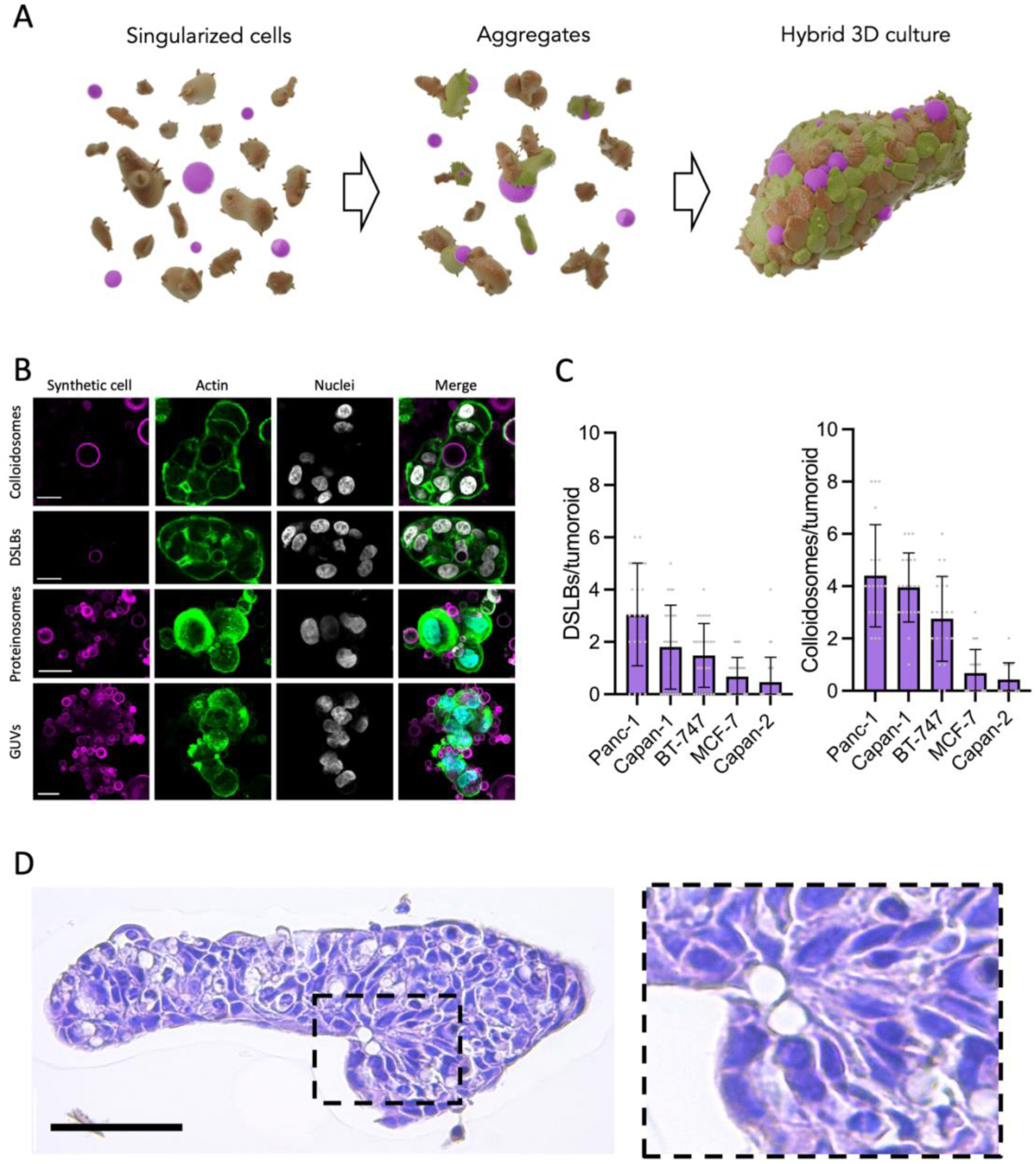
Self-assembly of hybrid tumoroids. **A**) Schematic illustration of the hybrid culture self-assembly process. Cancer cells (brown) and synthetic cells (magenta) are plated on low adhesion surfaces to induce aggregation and subsequent constriction of cell cluster to 3D cultures integrating synthetic cell models. **B**) Representative confocal microscopy images of Panc-1 tumoroids, stained for nuclei and actin, integrating synthetic cell models, 48 hours after formation. Scale bars are 20 µm. **C**) Quantification of DSLB- and colloidosome-based synthetic cells integration into tumoroids formed from various cell lines after 48 hours of co-culture. Results shown as mean ±SD from n>14 3D cultures per condition. **D)** Representative hematoxylin and eosin-stained histological slides of a Panc-1 tumoroid self-assembled with DSLB-based synthetic cells. Scale bar is 300 µm.

We screen several synthetic cell models together with tumoroid-forming cells to derive the biophysical principles underlaying synthetic cell-natural cell co-assembly. By equipping the synthetic cells with immuno-receptor ligands, we characterize the microstructural organization of signaling-competent interfaces between the synthetic and natural cells that allows for membrane-membrane protein signaling. Finally, we demonstrate the application of this approach by forming bottom-up assembled artificial tumor immune microenvironments (ART-TIMEs) within pancreatic ductal adenocarcinoma (PDAC) tumoroids to study immune receptor co-signaling along the PD-1 axis. With this, we probe PD-1/CD2 co-signaling as a driver for immune evasion. We envision this approach to yield a new generation of augmented 3D cultures with enhanced biomimetic capabilities for cancer research, but also to open new avenues for synthetic cell research, particularly to study interaction between living and non-living cells in a tissue-context.

## Results and Discussion

### Integration of synthetic cells into tumoroids by self-assembly

To assess the integration of synthetic cell models into tumoroids, we screened an initial panel of immortalized cancer cell lines (BT-747, MCF-7, Capan-1, Capan-2 and Panc-1) for self-assembly-based tumoroid formation from singularized cells by restricted adhesion. The cells were selected based on the expression of extra- and intercellular adhesion proteins (**Fig. S1A**), covering fast and slow growing cultures as well as representing high immunogenic (breast) and low immunogenic (PDAC) cancers. All cell lines formed tumoroids by self-assembly via restricted adhesion within 48 hours (**Fig. S1B**), whereby Panc-1 and BT-747 formed the most defined and condensed tumoroids. As synthetic cell candidates for integration and hybrid tumoroid formation, we produced a panel of routinely applied synthetic cells models, including GUVs^15^, polymer coacervates^10^, inorganic silica colloidosomes^13^, DSLBs^14^ and proteinosomes^12^. Upon co-incubation with singularized Panc-1 cells, we observed various levels of integration and attachment, where colloidosomes and DSLBs showed full integration into the self-assembled tumoroids and GUVs and proteinosomes showed strong attachment and partial integration (**Fig. 1B**). Coacervates did not integrate, even after prolonged incubation periods. Quantification of the number of integrated DSLB- and colloidosome-based synthetic cells per tumoroid for all cell lines revealed the highest integration rates in Panc-1 and Capan-1, followed by BT-747 tumoroids (**Fig. 1C**). Increasing the synthetic-to-natural cell ratio resulted in a proportional increase of integrated synthetic cells (**Fig. S2A**). This indicates that the integration mechanism is stochastic in nature, robust and not perturbating the self-assembly process of the cancer cells in a significant manner. Importantly, we observed retention of colloidosomes and DSLB within the tumoroid for a 7-day growth period, indicating stable incorporation and retention within the 3D architecture (**Fig. S2B**). The stable integration of the synthetic cells into the tissue micro-anatomy was also evident from histological analysis, where synthetic cells were embedded within the multicellular structure without larger surrounding ECM but with direct contact to the cancer cells (**Fig. 1D**). Taken together, these findings demonstrate that synthetic cells can participate in the self-assembly processes of living cells and stably integrate into tumoroids assemblies over long time periods. As the different synthetic cell models showed varying levels of integration, we further focused on resolving the biophysical mechanism for the self-assembly of these hybrids.

### Mechanism of hybrid-tumoroid assembly and the synthetic-natural cell interface

To assess the mechanism of synthetic cell integration, we first performed live-cell imaging of DSLBs incubated with singularized Panc-1 cells during tumoroid formation under restricted-adhesion conditions. We observed a simultaneous process in which cancer cells formed intercellular adhesions and aggregates, while also adhering to the synthetic cells, pulling them into the developing tumoroids (**Movie S1**). There was no indication for a sequential process, where initial tumoroid stages would form first and then integrate the synthetic cells. Moreover, integration of synthetic cells into pre-formed tumoroids was significantly limited (**Fig. S3A**), suggesting a process where adherence of cancer cells to synthetic cells, akin to their interaction with other adjacent cancer cells is required to form hybrid tumoroids.

To assess a potential active and adhesion-mediated integration mechanism, we performed confocal microscopy imaging of the synthetic cell - natural cell interface during self-assembly. We found that natural cells formed concentric membrane interfaces with the synthetic cells (**Fig. 2A**). These formed from actin-rich protrusions with a filopodia-like morphology, indicative of an extracellular adhesive contact sites. This view was further supported by transmission electron microscopy (TEM) analysis, where cellular projections and close membrane-membrane interactions between the natural cell and synthetic cells were observed (**Fig. 2B**). This suggests a mechanism of hybrid-tumoroid self-assembly driven by cellular adhesion to the synthetic cells. Importantly, the synthetic cells applied in these experiments did not feature specific biofunctionalization or adhesive ligands on their surface. However, the synthetic cells underwent significant opsonization with serum proteins, potentially mediating adhesion^16^ (**Fig. S3B**), which were also visible on the synthetic cell surface in the TEM analysis. Of note, the level of serum opsonization (i.e. the total amount of protein adsorbed) to the synthetic cells correlated with the degree of synthetic cell integration, where colloidosomes and DSLBs formed the largest protein coat, followed by GUVs and proteinosomes, while coacervates only had minimal opsonization (**Fig. 2C**). As the different cell lines integrated synthetic cells with varying efficacy, we hypothesized that hybrid tumoroid formation is directed by the balance of extra- and intercellular adhesion in the system, similarly to the formation of pure 3D cultures itself^17^. To test this hypothesis, we correlated the degree of synthetic cell integration with the expression of inter- and extracellular adhesion proteins by the different cell lines. We calculated the ratio of e-cadherin (intercellular adhesion) expression to expression of various integrin subunits (extracellular adhesion) from normalized transcripts per million data (see **Fig. S1A**). This approximation of the inter- and extracellular adhesion balance indicated that cells relaying more on extracellular adhesion (i.e. lower cadherin expression) also integrate more synthetic cells, most likely due to stronger interactions with their opsonized protein coat. In line with this, when DSLBs were coated with a passivating polyethylene glycol (PEG) layer coupled to the phospholipid head groups in their membrane, synthetic cell integration was significantly reduced (**Fig. S3C**).

**Figure 2.**
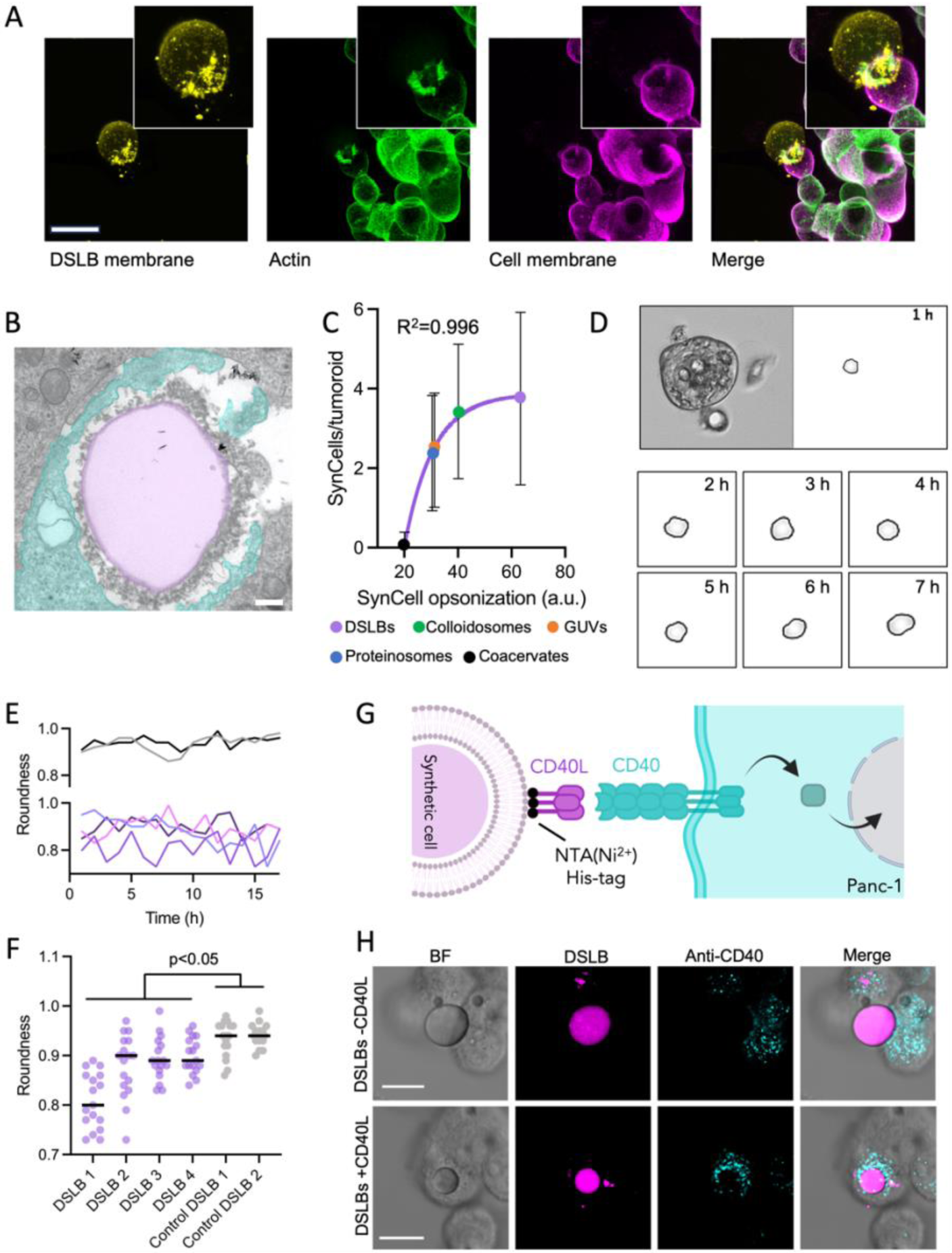
Adhesion-mediated self-assembly of hybrid tumoroids. **A)** Representative confocal microscopy maximal z-projection of a DSLB synthetic cell in the process of integrating into a Panc-1 tumoroid. Forming 3D aggregates were fixed 8 hours after co-incubation and stained for membrane (wheat germ agglutinin) and actin (phalloidin) to resolved adhesive interfaces between synthetic and natural cells. Inset shows magnified area from the synthetic cell contact site. Scale bar is 20 µm. **B)** Representative transmission electron microscopy image of a DSLB (magenta) integrated into a Panc-1 tumoroid showing cellular protrusion (cyan) around the synthetic cell. Scale bar is 500 nm. **C)** Correlation of synthetic cell serum opsonization with synthetic cell integration into Panc-1 tumoroids. Exponential fit is shown in magenta. Results shown as mean ±SD from n>14 tumoroids per condition. **D)** Bright field live cell imaging of a Panc-1 tumoroid with integrated DSLBs. An exemplary synthetic cell is segmented to derive deformation and roundness over time. **E) and F)** Quantification of roundness of four synthetic cells from sequences in D over 17 hours of observation (magenta). Two control synthetic cells sitting outside of the tumoroid (grey) shown as control. P value calculated by one-way ANOVA. **G)** Schematic illustration of the recombinant protein coupling to DSLB-based synthetic cells to induce receptor-ligand interactions with natural cells in tumoroids. **H)** Representative confocal microscopy maximal z-projections of DSLBs integrated into Panc-1 tumoroids without biofunctionalization (top panel) or with conjugated recombinant CD40L ectodomains (bottom panel). CD40 receptor localization in Panc-1 cells was detected by anti-CD40 staining after permeabilization. Scale bar is 10 µm.

Considering that hybrid tumoroid formation depends on an adsorbed adhesive interface, the presence of a sufficiently high interface tension on the synthetic cell surface would be required. GUVs, proteinosomes, DSLBs and colloidosomes have reported interface tension between 3 – 50 mN/m in physiological buffers^18–21^. In contrast, coacervates exhibited significantly lower interface tension (10 – 500 µN/m)^22,23^, consistent with their decreased opsonization and tumoroid integration. Importantly, the interactions between natural and synthetic cells were not merely transient and persisted post-integration. Time-resolved segmentation of individual synthetic cells within tumoroids from live cell imaging data revealed that elastic DSLBs, which have comparable stiffness to natural cells (∼1 kPa)^14^, undergo constant deformation within the tumoroids (**Fig. 2E** and **Movie S2**). This implies that natural cells maintain the adhesive interface to synthetic cells within the tumoroids and continuously exerting forces on them.

We next focused on specific biofunctionalization of the interface. Synthetic cells models can be readily functionalized with recombinant proteins to engage in receptor-ligand signaling with natural cells in *trans*^9^. This can effectively mimic the membrane-membrane interactions of cancer and immune cells in the TIME and adds a biochemical communication axis to the biomechanical interactions in the synthetic-natural cell interplay. To demonstrate this concept, we integrated NTA(Ni^2+^)-modified DSLBs presenting signaling-active recombinant polyhistidine-tagged CD40L ectodomains, a central regulatory protein on CD4^+^ T cell in the TIME^24^, into Panc-1 tumoroids and subsequently assessed CD40-mediated signaling within the cancer cells. Immuno-staining showed localization of CD40-enriched granule, similar to those observed between T cells and dendritic cells^25^, at the tumor-synthetic cell interface (**Fig. 3F**) and quantitative PCR analysis revealed increased transcription of the anti-apoptotic proteins Bfl-1, a known CD40 down-stream target gene expressed in Panc-1 cells^26,27^ (**Fig. S3D**). This demonstrates that synthetic cells within hybrid tumoroids can form receptor-ligand interfaces with cancer cells, triggering biomimetic down-stream signaling.

**Figure 3.**
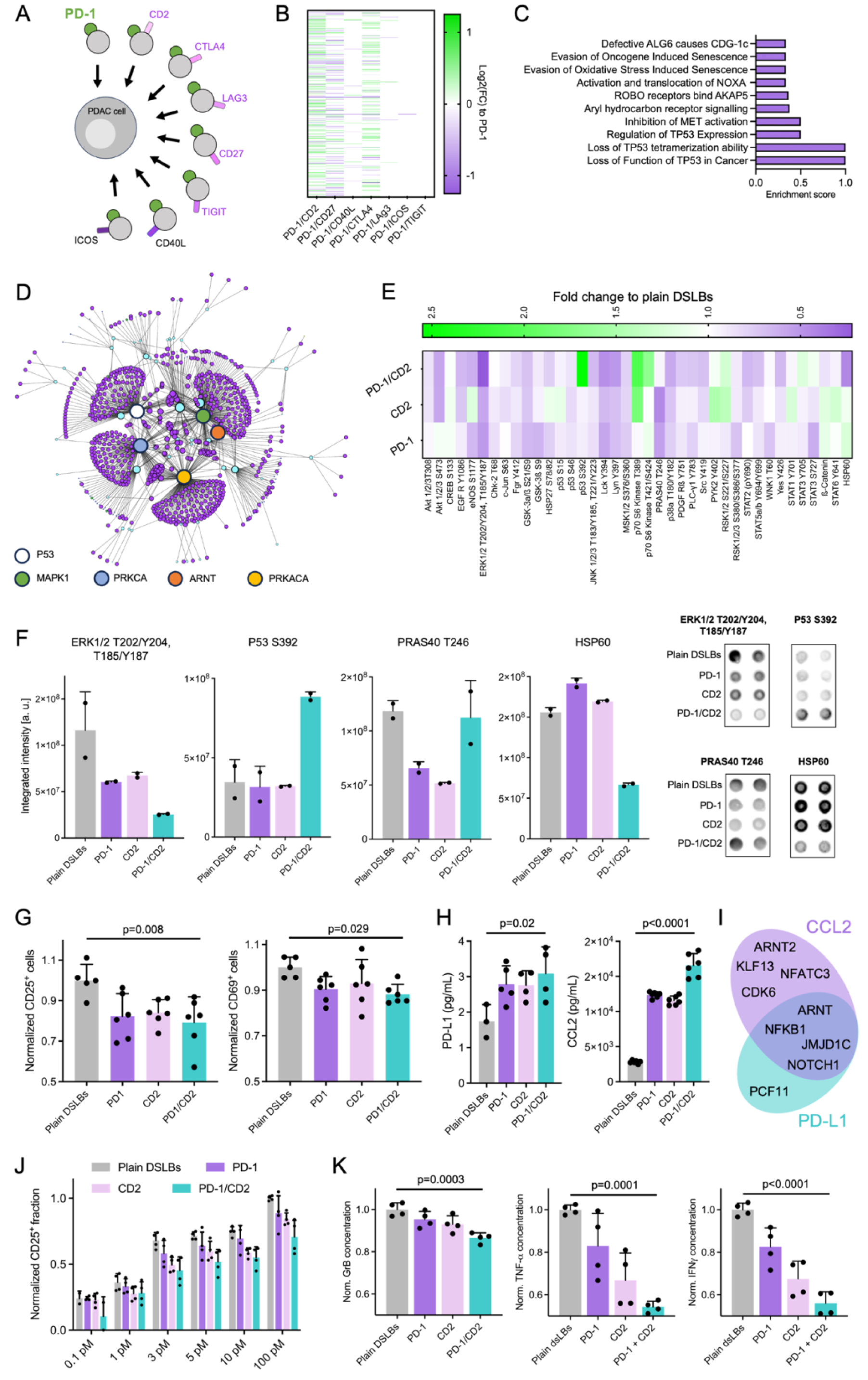
Artificial tumor immune microenvironments. **A)** Schematic representation of the ART-TIME screening approach and the panel of immune ligands tested. **B)** Differential gene expression analysis of Panc-1 tumoroids with integrated ART-TIMEs presenting varying combinations of immune ligands. DEGs compared to ART-TIMEs with PD-1 only are shown to derives transcriptional changes based on receptor co-signaling. **C)** Pathway enrichment analysis from PD-1/CD2 DEGs in C. Pathways with the ten highest enrichments scores are shown. **D)** Gene network analysis from PD-1/CD2 DEGs. 5 nodes with the highest degrees are highlighted. **E)** Heat map of protein phosphorylation analysis from Panc-1 tumoroids with ART-TIMEs of PD-1, CD2 and PD-1/CD2. Phosphorylation levels are shown as compared to hybrid tumoroid with plain DSLBs. **F)** Four exemplary protein phosphorylation levels from E, showing synergistic, paradox or *de novo* effect with PD-1/CD2 co-presentation. Results shown as mean from two antibody spots pooled from two replicates. **G)** Flow cytometry quantification of the fraction of CD25 and CD69 expressing primary human CD8 T cells incubated with pre-conditioned medium of Panc-1 tumoroids with varying ART-TIME compositions. Results are shown from n=3 donors as mean ±SD from two replicates each. **H)** Enzyme-linked immunosorbent assay-based quantification of soluble PD-L1 and CCL2 secreted from Panc-1 tumoroids with ART-TIMEs of varying composition after 48 hours of incubation. Results shown as mean ±SD from 2 biological replicates measured in duplicates. **I)** Ven diagram of PD-L1 and CCL2 transcription factors identified as DEGs in PD-1/CD2 ART-TIME tumoroids. **J)** Flow cytometry quantification of the fraction of CD25 expressing primary human CD8 T cells incubated with ART-TIME containing Panc-1 3D cultures for 12 hours in the presence of increasing bispecific T cell engager concentrations. Results are shown from n=2 donors as mean ±SD from two replicates each. **K)** Enzyme-linked immunosorbent assay-based quantification of three effector cytokines secreted from primary human CD8 T cells incubated with Panc-1 tumoroids with ART-TIMEs of varying composition after 12 hours of incubation with 5 pM bispecific T cell engager concentration. Results shown as mean ±SD from 2 biological replicates measured in duplicates. p values are indicated from one-way ANOVA statistical testing.

Taking these data together, a hybrid tumoroid formation model emerges, where synthetic cells provide an adhesive interface that natural cells use to equilibrate their inter- and extracellular adhesion balance during self-assembly and organization. This is in high similarity to the processes found during cellular self-assembly *in vivo*, where extra- and intercellular adhesion forces direct tissue organization^28^. As a key principle for synthetic cell engineering, our integration analysis reveals that for stable and biomimetic self-assembly of synthetic cells into 3D hybrids, the biophysical characteristics of synthetic cells, including their interface tension, protein opsonization, and elastic modulus needs to be considered along the natural cells inter-to extracellular adhesion balance. While specific conjugation of the synthetic cells with adhesive ligands is not required for integration, the hybrid interfaces within the tumoroids can be augmented with specific biomolecular interactions, providing defined molecular input signals and inducing receptor-signaling in the natural cells. Stable and long-term embodiment of these synthetic cells into tumoroids, opens the door for bottom-up reconstitution of an artificial TIME (ART-TIME) to systematically probe molecular interactions and specific receptor-ligand signaling axis in the cancer-immune interplay.

### Artificial tumor immune microenvironments to study receptor co-signaling in 3D

Having established that synthetic cells can co-assemble with living cells into hybrid tumoroids, we next aimed to demonstrate a specific application scenario where the synthetic cells can replace other cell types for studies of the tumor microenvironment. The appealing advantage of an ART-TIME, in contrast to conventional approaches to study signaling processes in 3D, is that biomolecular interactions can be probed in a defined and isolated setting of reduced molecular complexity, where interfering cellular processes or molecular interactions are deliberately excluded. Applying this principle, we systematically investigated PD-L1 reverse signaling in combination with other immune receptors (co-signaling), to understand their contribution to immune evasion in pancreatic ductal adenocarcinoma. Previous work has shown that activation of PD-1 on T cells *via* cancer-cell presented PD-L1 drives T cell suppression while also inducing adaptive PD-L1 reverse signaling into the cancer cells^29^. Several other receptors have been suggested to induce reverse signaling and co-localization with PD-1 such as CD2, CTLA4 or CD27^30–33^. However, the complexity of natural cells has hindered obtaining a functional mechanism and causality for such interactions.

To infer the relevance of co-reverse signaling of PD-L1 with ART-TIME, we first compiled a panel of potential co-signaling ligands with reported immuno-modulatory function presented on tumor infiltrating lymphocytes (**Fig. 3A**). Signaling-active ectodomains of these ligands were immobilized by polyhistidine-tag chemistry on the DSLB surface at biomimetic membrane densities (300 molecules/µm^2^) and self-assembled into Panc-1 tumoroids to form ART-TIMEs. The Panc-1 to synthetic cell ratio was adjust to 10:1 to emulate immune cell infiltration levels found in PDAC^34^. We first derived transcriptional changes in ART-TIME cultures by RNA-sequencing and differential gene expression analysis. Differentially expressed genes (DEGs) were derived from pair-wise comparisons between 3D hybrids with PD-1 DSLBs or with DSLBs co-presenting PD-1 with any of the ligands of the panel (**Fig. 3B and supplementary data**). No significant DEGs, hence no co-signaling, were found when TIGIT or ICOS were co-presented with PD-1 and only a limited number of DEGs was induced by additional presentation of CD40L and LAG3. However, co-presentation of PD-1 with CD27, CTLA4, and specifically CD2, induced a large number of DEGs. From these, pathway enrichment analysis revealed that specifically components of the p53 and AhR receptor signaling pathways were differentially regulated by co-signaling (**Fig. 3C and supplementary data**).

Based on reported indication that co-activation of PD-L1 and CD58 could be central for cancer immune evasion^33,35^, we further focused on their respective receptors, PD-1/CD2, and derived gene regulatory networks for these ART-TIMEs (**Fig. 3D and supplementary data**). Five central signaling pathways were represented in the networks with degrees >10, Mitogen-activated protein kinase 1 (MAPK1), protein kinase C alpha (PRKCA), protein kinase A (PRKACA), p53 and aryl hydrocarbon nuclear translocator (ARNT), suggesting larger scale adaptations of the cancer cells by co-signaling. We experimentally verified differential activation of the identified signaling kinases and transcription factors in the networks, by targeted phospho-kinase profiling (**Fig. 3E**). Differential phosphorylation was observed for several proteins, covering the five central signaling axis represented in the gene network analysis. Importantly, differential effects with synergistic, paradox or *de novo* phosphorylation were observed between PD1, CD2 and PD1/CD2 ART-TIMEs (**Fig. 3F**), highlighting the complex intracellular interplay induced by co-reverse signaling. Taken together, this suggests a functional mechanism where PD1 and CD2 together induce reverse signaling along four major axis and demonstrates how synthetic cells can be applied as bionics in tumoroids to emulate cancer immune interactions.

To resolve if PD-1/CD2-induced reverse signaling has relevant functional effects on the immuno-suppressive phenotype of PDAC tumoroids cultured with ART-TIMEs, we assessed the potential to these hybrid tumoroids to suppress T cell stimulation. We found that activation of primary human T cells was minorly but significantly reduced when they were cultured with preconditioned media from PD-1/CD2 ART-TIME tumoroids (**Fig. 3G**). This indicates that PD1/CD2 co-signaling promotes secretion of soluble immune suppressive factors from PDAC cells. In the preconditioned media, we identified elevated levels of CC-chemokine ligand 2 (CCL2) and the soluble ligand of PD-1 itself, PD-L1 (**Fig. 3H**). Both have been reported to mediate immune suppression in the TIME^36,37^. We linked the increased levels of these suppressors to our DEG analysis above, were several of their transcription factors showed increased expression, including ARNT (**Fig. 3I**), which constituted a central node in our gene regulatory network. Importantly, cross-signaling between AhR-ARNT with p53^38^ and with protein kinase C^39^ has been described previously, which links this cross-signaling to our differential phosphorylation analysis in the tumoroids with PD-1/CD2 ART-TIMEs (see **Fig. 3F**). Taken together, a signaling mechanism emerges where CD2 induces co-reverse signaling with PD-1 *via* p53 and protein kinases, which results in altered transcription involving AhR-ARNT to promote an immune suppressive PDAC phenotype.

To assess the relevance of this phenotypic adaptation in an immunotherapeutic context, we next incubated primary cytotoxic T cells with the PD-1/CD2 ART-TIME tumoroids in the presence of CD3/Her2-directed bispecific T cell engager antibody. T cell engager antibodies linking CD3 on T cells with tumor antigens of the human epidermal growth factor receptor family have demonstrated broad efficacy in clinical trials for PDAC^40^. This T cell engager-mediated recognition and killing of PDAC cells by T cells can be reproduce in *in vitro* cultures^14^. We observed dose-dependent killing in a 0.1-100 pM range of engagers but did not resolve any significant differences in their efficacy when comparing tumoroids containing plain DSLBs and PD-1/CD2 ART-TIMEs (**Fig. S4A**). However, when isolating the T cell from these experiments after incubation with the tumoroids and quantifying the expression of T cell activation markers CD25 and CD69, we observed significant differences, with lower activation of the T cells in PD-1/CD2 ART-TIMEs (**Fig. 3J and Fig. S4B**). Moreover, when quantifying the levels of pro-inflammatory and effector cytokines secreted from the T cells, we found significantly lower secretion in PD-1/CD2 ART-TIMEs (**Fig. 3K**). Taken together, this demonstrates that presentation of PD-1/CD2 induces adaptive downstream signaling in PDAC tumoroids, leading to a more immuno-suppressive phenotype that suppresses T cell activation in bispecific T cell engager-mediated killing.

Finally, to support the finding that PD-1/CD2 co-signaling could impact on PDAC immune interactions *in vivo*, we verified the relevance of this co-signaling axis for clinical outcomes of PDAC patients. We found significant co-expression of the receptors (CD2/PD-1) and ligands (CD58/PD-L1) in cancerous and non-transformed tissue samples from transcriptomic data from a cohort of 176 patients tumor samples and 252 normal samples (**Fig. S5A**). By dividing the patient cohort in CD58/PD-L1^high^ and CD58/PD-L1^low^ populations, we found that co-expression of the PD-1/CD2 ligands is indeed correlated to overall survival (**Fig. S5B**). This supports the notion that PD-1/CD2 co-signaling has clinically relevant effects on the interplay between PDAC and the host immune system.

## Conclusion

In conclusion, our study provides compelling evidence that synthetic cell models can be stably integrated into 3D cultures through adhesion processes during cellular self-assembly. This integration enables systematic screening of microenvironmental factors within tumoroids.

Our initial screening, which evaluated seven receptor ligands for co-signaling, demonstrates the feasibility of this approach. Future studies could upscale this screening to assemble more complete TIMEs in a step-by-step approach towards reconstituting more complete replicas of tumor infiltrating leukocytes within the tumoroids.

However, already the minimalistic ART-TIME model we developed, shows a significant impact on cancer cell phenotypes, as evidenced by the suppression observed in our supernatant assays. We elucidate the functional mechanism of PD-1/CD2 co-signaling, which induces protein kinase- and p53-mediated downstream signaling, leading to the secretion of soluble suppressive factors regulated by transcription factors such as ARNT. This highlights the plasticity of the TIME and how cancer cells sense and respond in a contact-dependent manner to the presence of cellular TIME components. Our approach holds potential for high-throughput, *ex vivo* drug screening, allowing for the step-by-step assembly and assessment of individual signaling axes within a biomimetic, biophysically adjusted context that not only mirrors the surface receptor density of cells in the TIME but also their stiffness and membrane mobility. While the *in vivo* TIME is composed of heterogeneous cell types, with varying surface receptor expression that is currently not replicated in full detail in ART-TIMEs, they provide a valid framework to evaluate the contributions of specific signaling pathways. By this, signaling analyses, which could not be performed in 3D tumoroids otherwise, can isolate interactions and systematically probe the effects of co-signaling.

Furthermore, our study underscores the advancement of synthetic cell engineering, showcasing several synthetic cell models that can functionally merge with living matter. We identify adhesion as a central factor in coupling synthetic cells with the self-assembly properties of living cells. But not only biochemical signaling is crucial for integration, also the mechanical properties of the synthetic cells are important, as we observed significant deformation of the DSLBs once integrated into the tumoroids. Our synthetic cell screening serves as a blueprint to guide further optimization and functional integration of synthetic and living cells. The fact that we observed stable incorporation of the synthetic cells within tumoroids for several days, is indicative for a true integration where the synthetic cells are accepted into the multicellular collective and unify into a coherent tissue architecture. Overall, our study marks a significant step towards the functional integration of living and non-living matter, advancing the field of synthetic cell engineering and its application in biomedicine.

## Acknowledgments

We would like to thank the INM Fluorescence Microscopy Core Facility, Cao Nguyen Duong and Angela Rutz (Leibniz Institute for New Materials), Andrea Hellwig (Neurobiology, Heidelberg University) for help in experiment design and technical support. The authors acknowledge funding from the Pharmazeutische Forschungsallianz Saarland, the Daimler and Benz Foundation (32-12/22), the Joachim Herz Foundation (Add-on Fellowship and Innovate! Akademie), German Science Foundation (Emmy Noether Program, project number 525255627 and 545610076), and the ERC Marie Sklodowska Curie Actions/UK Engineering and Physical Science Research Council (E. Pantano/X023907/1).

## Materials and Methods

### Confocal microscopy

Confocal microscopy was conducted using an LSM 880 laser scanning microscope (Carl Zeiss AG). Images were captured using a 63× immersion oil objective (Plan-Apochromat 63x/1.4 oil DIC M27, Carl Zeiss AG, Germany). The images were analyzed using ImageJ software. For imaging, samples were transferred to 8-well Nunc LabTeK glass bottom chamber slides filled with 200 µL PBS.

### DSLB assembly

DsLBs were assembled following a strategy similar to our previous protocol^14^. Briefly, 100 mg of PDMS (Sylgard 184, Dow Corning USA) were mixed with 750 µL PBS supplemented with 1 mM SDS and crudely emulsified by resuspension. The solution was then incubated in a sonication bath at room temperature for 2 min to produce a dispersed o/w emulsion. Thereafter, MgCl2 (Sigma Aldrich, Germany) at a final concentration of 40 mM was co-added with 200 µL of a 6 mM (final lipid concentration) SUV solution to generate DSLBs. The SUVs, composed of 74 mol% EggPC, 20 mol% EggPG, 5 mol% PE-DGS-NTA(Ni^2+^) and 1 mol% LissRhodamine B-PE or Atto488-PE (all Avanti Polar Lipids, USA) were produced by extrusion. For experiments involving PEGylated DSLBs, PEG750 head group-modified 16:0 PE lipids were added to a final concentration of 10 mol% to the SUV mixture. The DSLBsolution was incubated at room temperature, protected from light, for 2 min and subsequently pelleted by centrifugation at 8600 xg for 30 sec. To wash the DSLBs, the supernatant was discarded, and the pellet was resuspended in 1 mL PBS, followed by a centrifugation step with the same parameters specified above. This procedure was then repeated a second time. Subsequently, the DSLB solution was resuspended in 1 ml PBS, transferred to a clean vessel and stored light-protected at 4 °C until further use.

### Assembly of other synthetic cell models

For synthetic cell assembly, previously established protocols were adopted. Colloidosomes were formed from inorganic silica nanoparticles following the protocol described by Li et al.^13^. Proteinosomes were assembled from poly(*N*-isopropylacrylamide)-modified BSA nanoconjugates as described by Huang et al.^12^. For GUV assembly, a bulk emulsification approach and phase transfer method was applied as described by Staufer et al.^15^. Coacervates were formed from poly(diallyldimethylammonium chloride)/ adenosine triphosphate mixtures as described by Moreau et al.^41^.

### Recombinant protein coupling protocol

For the decoration of DSLBs with proteins, the total amount of accessible membrane in a DSLB production batch was calculated. This was achieved via calibration against an SUV dilution series prepared in a 96 well plate with defined concentration. The fluorescent intensity, originating from either the Rhodamine B or Atto 488 incorporated in the DSLB membrane, was measured with a TECAN Spark (Tecan Group, Switzerland) plate reader. The device was controlled with the TECAN SparkControl software, excitation/emission were adjusted to 537/582 nm for Rhodamine B and 495/595 nm for Atto 488, respectively. The total lipid concentration of a DSLB solution was quantified by measuring its fluorescence intensity in conjunction with an SUV dilution series, the latter being used to plot a calibration curve. The amount of total accessible DGS-NTA(Ni2+) was derived from the final lipid concentration. The recombinant proteins PD-1, CD2, CD27, CD40 Ligand, CTLA-4, LAG3, ICOS and TIGIT (all Sino Biological, China) were added to aliquots of a DSLB production batch at a molar ratio of 2:1 (protein to DGS-NTA(Ni2+)) either alone or as a blend of PD-1 and one additional protein. Both proteins in the latter case featured a molar ratio of 2:1 and were premixed. The DSLB-protein solution was incubated for 60 min at 4°C while protected from light and subsequently washed once by centrifugation and resuspension in PBS. The final DSLB lipid concentration was calculated via plate reader quantification as described above.

### Recombinant proteins, sources and accession numbers

| rec. proteins | source | catalogue nr. | NCBI ref. sequence |
| --- | --- | --- | --- |
| rec. hu. PD-1 (ECD, His Tag) | Sino Biological | 10377-H08H | NP_005009.2 |
| rec. hu. CD2 (ECD, His Tag) | Sino Biological | 10982-H08H | NP_001758.2 |
| rec. hu. CD27 (His Tag) | Sino Biological | 10039-H08B1 | NP_001233.2 |
| rec. hu. CD40 Ligand (His Tag) | Sino Biological | 10239-H08E | NP_000065.1 |
| rec. hu. CTLA-4 (His Tag) | Sino Biological | 11159-H08H | NP_005205.2 |
| rec. hu. LAG-3 (ECD, His Tag) | Sino Biological | 16498-H08H | NP_002277.4 |
| rec. hu. ICOS (His & hFc Tag) | Sino Biological | 10344-H03H | NP_036224.1 |
| rec. hu. TIGIT (ECD, His Tag) | Sino Biological | 10917-H08H-B | NP_776160.2 |

### Cell culture of Panc-1 culture, MCF-7 culture, Capan-1, Capan-2, BT-747

Cultivation of all cell lines was carried out in flasks treated for cell culture at 37°C, 5% CO2 and 100% humidity. The cell lines were grown in RPMI 1640 w/ L-Glutamine (VWR, Germany) medium supplemented with 1% penicillin/streptomycin (Gibco, Germany) and either 10% fetal bovine serum (Gibco, Germany) for Panc-1, MCF-7 and BT-747 or 20% fetal bovine serum for Capan-1 and Capan-2. Cells were split twice a week once they had reached a confluence of 80 to 90 %, at a ratio of 1:4 to 1:6. For approximation of normalized adhesion protein expression levels in different cell lines, transcripts per million data was accessed from the Human Protein Atlas under the given protein names.

### Hybrid assembly protocol

Cell lines (Panc-1 if not indicated otherwise) with a confluence of 80-90% were washed twice with PBS and detached from their respective flasks using Trypsin 0,25% - EDTA in HBSS (VWR, Germany). The cell suspension was washed with medium, pelleted and resuspended in RPMI 1640 w/ L-Glutamine medium supplemented with 1% penicillin/streptomycin and 1% fetal bovine serum. The cell concentration was determined via Neubauer improved counting chamber (Marienfeld, Germany) and adjusted to roughly 1*10^6 1/mL. To generate hybrid spheroids, 50.000 to 100.000 cells were mixed with untreated or protein-labelled DSLBs (utilizing a cell to DSLB ratio of 1 to 10 if not indicated otherwise) in untreated Nunc 96-Well Flat Bottom plates (Thermo Fisher Scientific, USA). Each well was adjusted to a final volume of 200µL using the spheroid culture medium described above. The spheroids were left to mature for 48 hours at 37°C, 5% CO2 and 100% humidity.

### Counting of SynCell/tumoroid

For analysis of synthetic cell integration efficacy into the various tumoroids, hybrid were formed as described above and culture for 48 hours. Hybrid cultures were subsequently washed by centrifugation with PBS and fixed for 15 min with 2% paraformaldehyde. Hybrid cultures were then image by bright field and fluorescence microscopy, recording the fluorescence signal from the various synthetic cell models. For GUVs and DSLBs, fluorophore-coupled lipids were integrated into the membrane, for proteinosomes and colloidosomes, 1% of AlexaFluor647-coupled BSA was added to the synthetic cells and for coacervates, 1% FITC sodium salt was dissolved into the PDDA phase. Single synthetic cells where then manually counted for each 3D culture.

### SDS-PAGE

To quantify the serum protein opsonization of the different synthetic cell models, each synthetic cell type was incubated with fully supplemented cell culture medium for 2 hours at 37°C at a final synthetic cell concentration of 500,000 cells/ mL. Subsequently, the synthetic cells were washed by gentle centrifugation or gravity sedimentation twice with PBS. The total protein concentration in the final samples was measured by bicinchoninic acid assays and run on a NuPAGE bold bis-tris 4 to 12% gradient gel with MES running buffer for a gel electrophoretic analysis. Electrophoresis was performed at 200 V for 35 min under denaturing conditions with a total of 3 μg loaded onto separate lanes. Protein staining was performed with Coomassie R250.

### Expression analysis of Bfl-1

To measure the expression levels of Bfl-1 in hybrid 3D cultures, the 3D cultures were collected after 49 hours of incubation. Total RNA was isolated using the RNeasy Mini Kit (Qiagen) according to the manufacturer’s instructions. A DNase digestion step (RNase-Free DNase Set, Qiagen) was included to eliminate any residual genomic DNA. The quality and concentration of the isolated RNA were assessed using a NanoDrop spectrophotometer (Thermo Fisher Scientific). cDNA was synthesized from 1 µg of total RNA using the High-Capacity cDNA Reverse Transcription Kit (Applied Biosystems) according to the manufacturer’s instructions. The reaction mixture consisted of random primers, dNTP mix, MultiScribe Reverse Transcriptase, RNase inhibitor, and buffer. Reverse transcription was carried out in a thermal cycler at 25°C for 10 minutes, 37°C for 120 minutes, and 85°C for 5 minutes. The synthesized cDNA was stored at −20°C until further use. The expression of Bfl-1 mRNA was quantified using SYBR Green-based qPCR. Reactions were prepared using PowerUp SYBR Green Master Mix (Applied Biosystems) in a final volume of 20 µL, containing 10 ng of cDNA, 0.5 µM of forward primer (5’-ACTGGAAGTGCGAGATGGAG-3’), 0.5 µM of reverse primer (5’-AGGGTGGTTGAGGTGACAGT-3’), and nuclease-free water. qPCR was performed in a QuantStudio 5 Real-Time PCR System (Applied Biosystems) under the following thermal cycling conditions: 95°C for 2 minutes (initial denaturation), followed by 40 cycles of 95°C for 15 seconds (denaturation) and 60°C for 1 minute (annealing/extension). A melt curve analysis was conducted to verify the specificity of the amplification. The relative expression levels of Bfl-1 mRNA were normalized to the expression of a housekeeping gene, GAPDH, using the ΔΔCt method. Each sample was run in triplicate, and negative controls without template (NTC) were included to confirm the absence of contamination. Data were analyzed using QuantStudio Design & Analysis Software (Applied Biosystems), and fold changes in gene expression were calculated relative to the controls.

### RNAseq, DEG analysis

For every condition, hybrid spheroids were generated in triplicates using 100.000 cells mixed with DSLBs at a ratio of 1 to 3 cells to DSLBs. DSLBs were coupled with either PD-1 alone or PD-1 and a second protein (CD2, CD27, CD40 Ligand, CTLA-4, LAG3, ICOS or TIGIT). DSLB-free spheroids and hybrid spheroids generated with uncoupled DSLBs served as negative controls. This set of conditions was created and processed in three parallel biological replicates. After the spheroid-formation had concluded, the 3D cultures were harvested and washed with ice-cold PBS. The supernatant was removed after a centrifugation step and the spheroid pellets snap-frozen in liquid nitrogen. RNAseq libraries were prepared using 150 ng of total RNA per sample with the NEBnext Ultra II Directional RNA library kit for Illumina (New England Biolabs (NEB); Ipswich, USA) according to the manufacturer’s recommendations. Libraries were quantified with the NEBnext library quant kit for Illumina (NEB) and then sequenced for approximately 40 – 60M 1x 75 nt single reads on a NextSeq500 (Illumina; San Diego, USA). After quality checks with FastQC Version 0.11.2 (http://www.bioin forma tics.bbsrc.ac.uk/proje cts/fastq c/), reads were demultiplexed with samtools (v1.3.1) and adaptor-trimmed (Q < 20) with Cutadapt (Version 1.4.132) using Trim Galore! (Version 0.3.3) (https://www.bioin forma tics.babra ham.ac.uk/proje cts/trim_galor e/). Reads were aligned to hg38 assembly with the grape-nf pipeline (https://github.com/guigo lab/grape -nf) wrapping STAR (Version 2.4.0j33) and RSEM (Version 1.2.2134). Pairwise differential comparisons were performed in R using the DESeq2 package.

### Pathway enrichment analysis and gene regulatory network analysis

Pathway enrichment data were obtained by analyzing DEG lists generated by comparing hybrid spheroid sequencing data with their negative controls. The Reactome website (https://reactome.org/) was used to identify significantly affected signaling pathways. An enrichment score was calculated by determining the ratio of impacted reactions to total reactions for the respective pathways. Gene regulatory networks were generated with the visual analytics platform NetworkAnalyst 3.0^42^.

### Survival and differential expression analysis

Pancreatic adenocarcinoma and normal pancreatic tissue gene expression data was used from TCGA (https://portal.gdc.cancer.gov/) and from GTEx (https://gtexportal.org). Gene expression data was DESeq2 normalized and scaled to have a mean expression of 1000 in each sample. Differential gene expression was determined using Mann-Whitney U-test. Survival analysis was performed using overall survival data and by using the gene expression to stratify patient samples. Survival analysis was performed by employing Cox proportional hazards regression. To comprehensively assess potential correlations across different expression thresholds, we examined all possible cutoff values between the 25th and 75th percentiles of gene expression. Multiple hypothesis testing was addressed by calculating the false discovery rate (FDR) using the Benjamini-Hochberg procedure. The optimal cutoff value was determined based on the highest statistical significance (lowest p-value and corresponding minimum FDR), and this cutoff was used in the Kaplan-Meier analysis to visualize survival differences.

### T cell isolation

Primary human CD8+ T cells were isolated from leukapheresis reduction system (LRS) chambers from de-identified, voluntary healthy blood donors utilizing the RosetteSep Human CD8+ T cell Enrichment Cocktail (STEMCELL technologies, Germany) negative selection kit following the manufacturers instruction. The cells were routinely cultivated in RPMI 1640 w/ L-Glutamine medium supplemented with 10% fetal bovine serum, 1% penicillin/streptomycin, 1% non-essential amino acids (Biowest, France) and 50 mM HEPES (Sigma Aldrich, Germany) in flasks treated for cell culture at 37°C, 5% CO2 and 100% humidity. For culture, expansion and all functional assays, the medium was further supplemented with 100 U/mL recombinant human IL-2 (STEMCELL technologies, Germany). Blood samples were provided by the Institute for Clinical Hemostaseology and Transfusion Medicine, Saarland University Medical Center, following ethics agreement number 34/23 (Ethikkommission Arztekammer des Saarlandes).

### T cell activation and expansion

Human CD8+ T cells were either cryopreserved immediately after isolation in a naïve state or activated and expanded. For this purpose, the cells were adjusted to a concentration of 1*10^6 1/mL and cultured together with Dynabeads human T-Activator CD3/CD28 beads (Gibco, Germany) according to the manufacturers instructions for 3 days. The beads were then separated from the cells by a magnetic field, while the T cells were supplemented with fresh medium, adjusted to a concentration of 1*10^6 1/mL and further cultured for 2 days. Finally, the T cells were cryopreserved until further use.

### T cell staining and flow cytometry

T cells were pelleted and resuspended in PBS containing 1% BSA (Sigma Aldrich, Germany) and transferred into U-bottom 96 well plates. Subsequently, anti-CD25 AlexaFluor488 (BC96, BioLegend, UK) and anti-CD69 AlexaFluor647 (FN50, BioLegend, UK) staining antibodies were added (1:400 in PBS +1% BSA). Samples were incubated at room temperature, light-protected for 30 min. Afterwards, the remaining unbound antibodies were removed by centrifugation and resuspension in PBS containing 1% BSA. The supernatant was removed after an additional centrifugation step and the cells were subsequently fixed in 2% PFA (Sigma Aldrich, Germany) in PBS for 30 min at room temperature under light-protection. A final centrifugation step was employed to replace the PFA, the cells were then resuspended in PBS containing 1% BSA and stored light-protected at 4°C until further use.

Staining of surface markers was quantified using an Attune NxT flow cytometer (Thermo Fisher Scientific, Germany) fitted with 488 nm and 637 nm laser lines. For every condition, a minimum of 10.000 T cells were analyzed. Data analysis was performed by the FlowJo V.10 software (FlowJo LLC, USA).

### Proteome profiler

Hybrid spheroids were generated using 100.000 cells mixed with DSLBs at a ratio of 1 to 8 cells to DSLBs. The DSLBs were decorated with either PD-1, CD2 or both proteins. Hybrid spheroids incorporating unlabeled DSLBs served as negative control. 40 technical replicates per condition were created in order to achieve a sufficient protein concentration. After the hybrid spheroids had fully formed, samples were washed with PBS and pooled. Subsequently, the spheroids were lyzed in Lysis Buffer 17 supplemented with 10 µg/mL Aprotinin, 10 µg/mL Leupeptin hemisulfate and 10 µg/mL Pepstatin A (all R&D Systems, USA) for 30 min at 4°C under constant shaking. Cell debris was removed via centrifugation and the protein concentration in the supernatant determined utilizing the BCA-Protein-Assaykit (Merck, Germany) according to the manufacturers instructions. From each condition, 600 µg of protein was used to load the membranes of the Proteome Profiler Human Phospho-Kinase Array Kit (R&D Systems, USA). The Assay was performed according to the specifications provided by the manufacturer. The Streptavidin-HRP included in the kit was substituted with IRDye 800CW Streptavidin (LI-COR Biotechnology, USA). Fluorescence intensity of the stained membranes was documented in an ODYSSEY M (LI-COR Biotechnology, USA) imaging system equipped with a 785 nm laser line. The images were analyzed using ImageJ (NIH) software.

### Bispecific T cell engager assays Assays

Hybrid spheroids were generated using 50.000 cells mixed with DSLBs (ratio 1:10) decorated with either PD-1, CD2 or both proteins. As a negative control, hybrid spheroids incorporating unlabeled DSLBs were utilized. After the hybrid spheroids had fully matured, the culture medium was replaced by RPMI 1640 w/ L-Glutamine medium supplemented with 1% fetal bovine serum, 1% penicillin/streptomycin, 1% non-essential amino acids, 50 mM HEPES and 100 U/mL recombinant human IL-2. A dilution series of an anti-Her2-CD3 BiTE antibody [XXX] was added to the samples, whereby the concentrations used are indicated in the figure legends. T cells activated as described above were removed from cryopreservation and added to the spheroid-antibody preparations. A total of 200.000 T cells per well were added (a ratio of Panc-1 cells to T cells of 1 to 2). Two distinct T cell donors, each in technical duplicates, were used to account for donor variation. After 12 to 48 hours of co-cultivation at 37°C, 5% CO2 and 100% humidity, the assay was terminated and the cells and supernatant harvested for further experiments.

### LDH Assays

To quantify T cell-mediated cytotoxicity in the BiTE assays, an LDH-Cytox Assay Kit (BioLegend, UK) was used. Hybrid spheroids that were cultivated without T cells and BiTEs were mixed with 40 µL of the provided lysis buffer and incubated for 1h at 37°C. The samples were resuspended every 15 minutes and served as a positive control. Subsequently, the BiTE assays were pelleted via centrifugation and 50 µL supernatant transferred to new 96 well plates. The LDH assay was performed according to the instructions provided by the manufacturer. The colorimetric reaction was conducted under light protection at room temperature for 15 to 20 min. Absorbance at 490 nm was measured at a TECAN Spark plate reader.

### ELLA assays

#### Tumor-associated proteins

Hybrid spheroids were generated using 100.000 Panc-1 or MCF-7 cells mixed with DSLBs at a ratio of 1:10. The DSLBs were decorated with either PD-1 alone, CD2 alone or PD-1 mixed with a second protein (CD2, CD27, CD40 Ligand, CTLA-4, LAG3, ICOS or TIGIT). Spheroids were left to mature for 48 hours, afterwards supernatant was harvested from all conditions. For the quantification of protein concentrations in the supernatant, two Custom Simple Plex Assay cartridges (Biotechne, USA) were employed according to the instructions provided by the manufacturer. Cartridge A was designed for the quantification of PD-L1, CCL2 and CTLA-4, while cartridge B tested for Fas, LAG3, MMP-9 and THBS. For cartridge A, half of the harvested supernatants were diluted with an equal volume of the sample diluent SD13 provided, whereas for cartridge B the samples were diluted with a fivefold excess of diluent. Quantification of protein concentrations was performed in the Ella Automated ELISA (Biotechne, USA) system.

#### T cell-associated cytokines

A BiTE Assay was performed as described above. After termination, supernatant was collected from all conditions and stored at −80°C. Subsequently, the samples were used to load a Custom Simple Plex Assay cartridge according to the manufacturers instructions. The cartridge quantified the concentration of the following proteins: IL-4, IL-6, IL-10, IL-17A, CCL19, IFN-γ, Granzyme B and TNF-α. Supernatant samples were mixed with an equal volume of the provided sample diluent SD13. Quantification of protein concentrations was performed in the Ella Automated ELISA system.

### Hybrid culture supernatant assays with T cells

Hybrid Spheroids were generated in the same fashion as described for the BiTE Assays. Once fully matured, supernatant from the different conditions was harvested. Naïve and activated T cells, stimulated as outlined above, were seeded in a 96 well plate with a density of 40.000 cells per well. In total, 2 donors of naïve T cells and 2 donors of activated T cells were used for the assay, each condition was prepared in technical duplicates. Dynabeads human T-Activator CD3/CD28 beads were mixed with the cells according to the manufacturers instructions. Each well was loaded with either 10 µL or 50 µL of the harvested hybrid spheroid supernatant and the final volume adjusted to 200 µL. Supernatant-free cultures and T cells supplemented with 55,6 ng/mL CD2 and 75,6 ng/mL PD-1 (0.1% of the concentration used for DSLB labeling) served as controls. Cultivation was terminated after 48 hours and the cells were stained for activation surface markers and fixed as described above.

