## Supplementary Information and Data for "Self-assembly of hybrid 3D cultures by integrating living and synthetic cells"

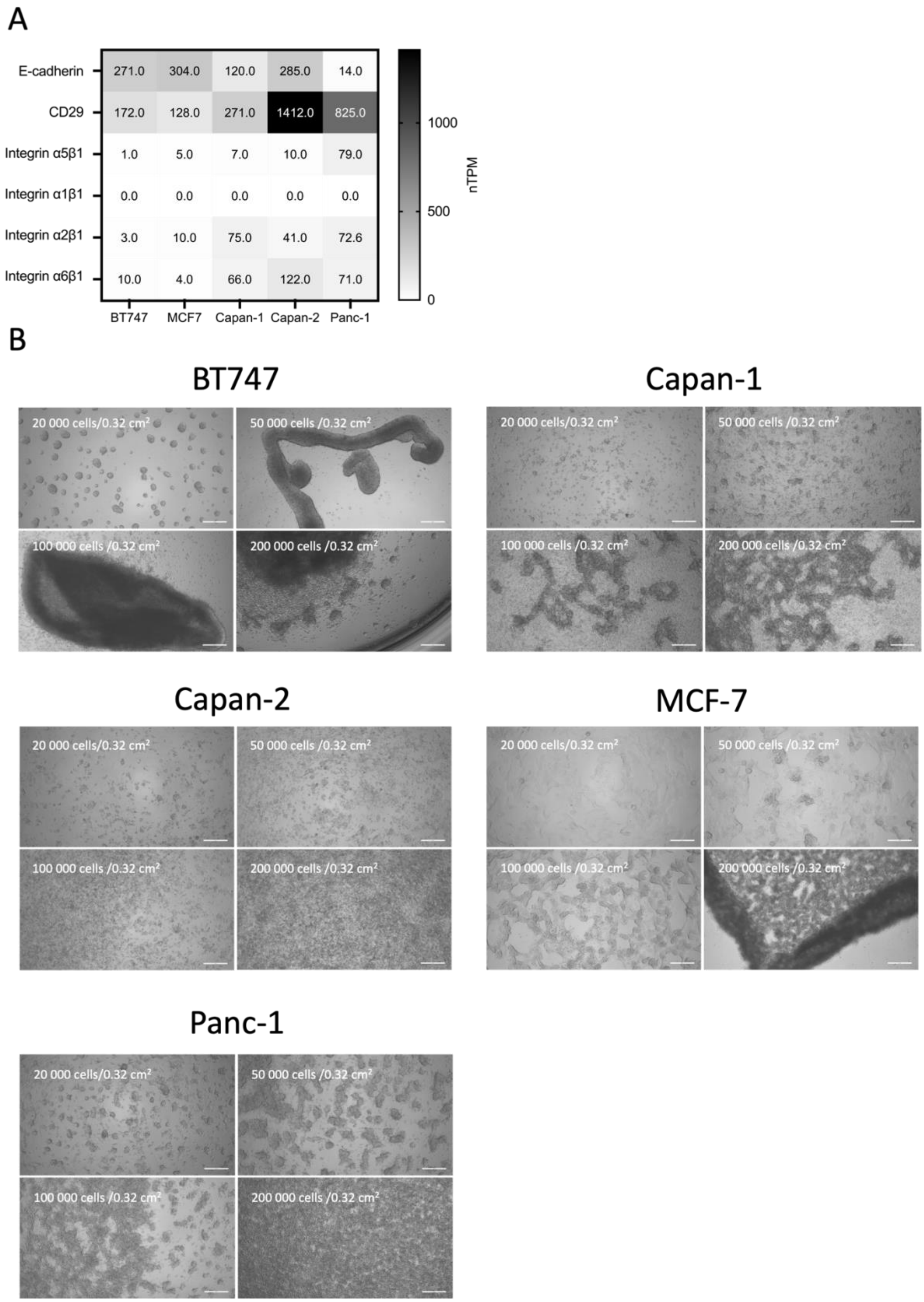


**Figure S1:** 3D cell culture cell lines. **A**) Normalized transcripts per million data for selected intercellular (cadherin) and extracellular (integrins) adhesion proteins in of the cell lines used in this study. Data was derived from the human protein atlas (proteinatlas.org). **B**) Brigth field microscopy data of the five cell lines used in this study, plated a various densities on low adhsive surfaces to induce 3D culture formation. Scale bars are 100 µm.

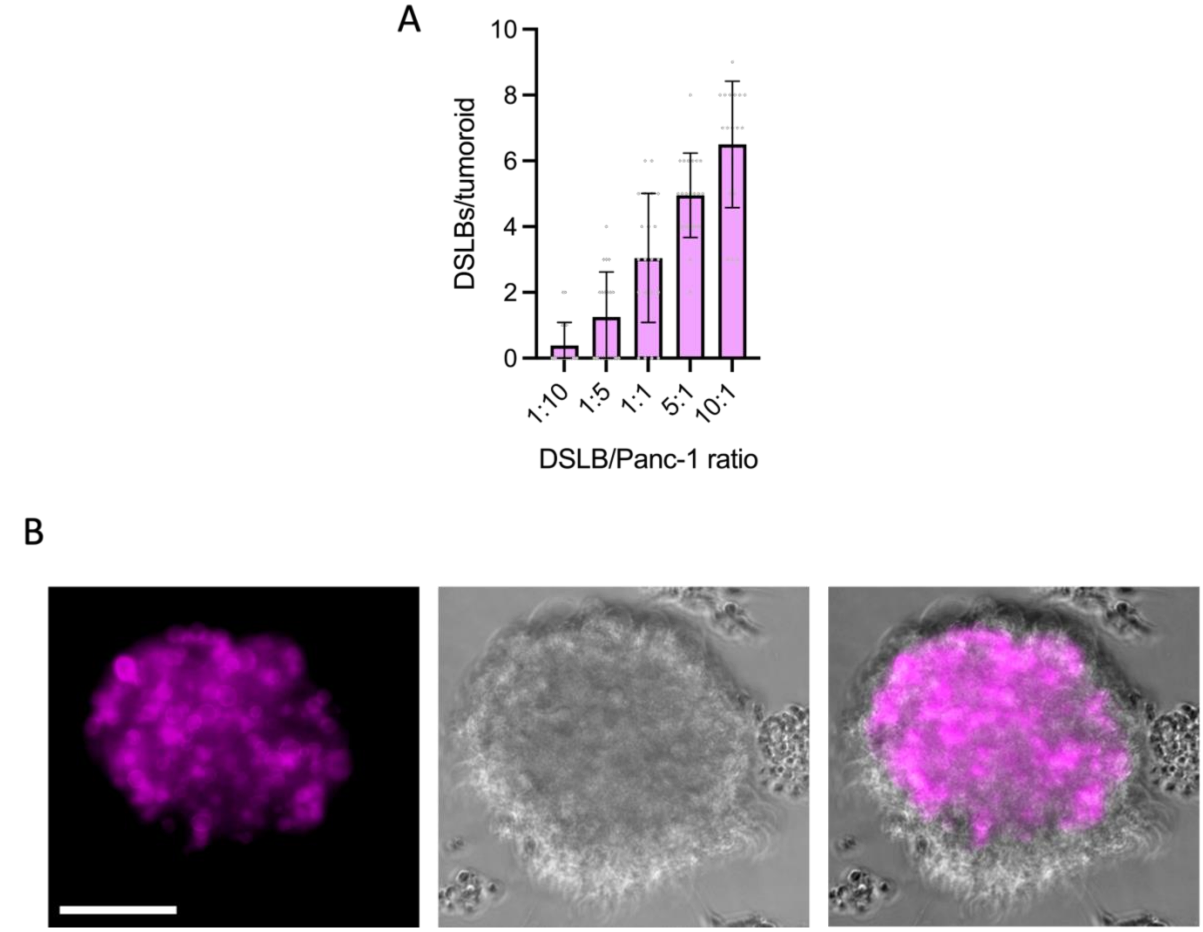


**Figure S2**: Stable integration of synthetic cells into tumoroids. **A)** Quantification of DSLB-based synthetic cells integration into 3D cultures formed from Panc-1 cells after 48 hours of co-culture in various synthetic cell to Panc-1 ratios. Results shown as mean ±SD from n>21. **B)** Representative confocal microscopy maximal z-projections of DSLB based synthetic cells integrated into Panc-1 tumoroids 7 days after formation. Scale bar is 200 µm.

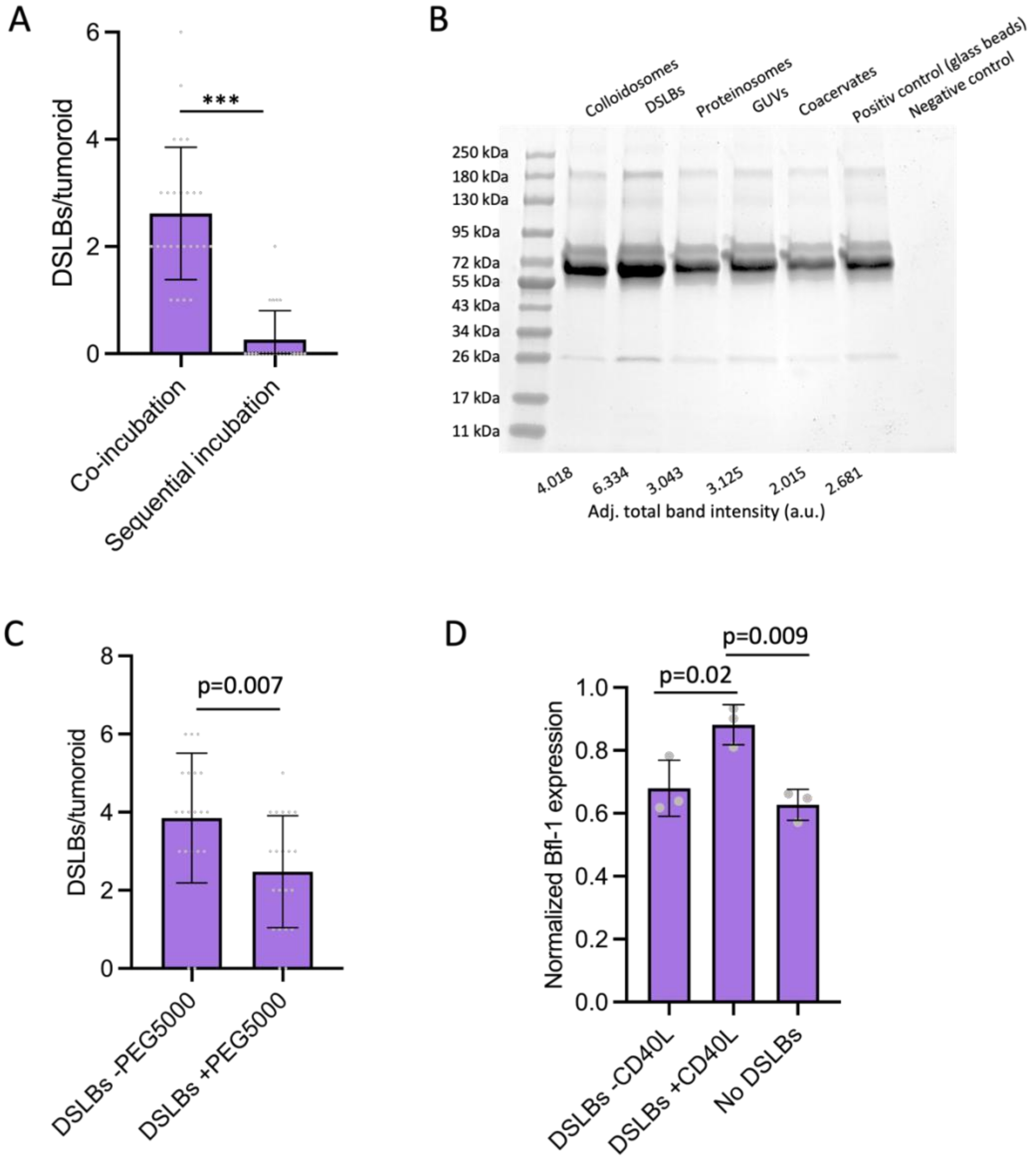


**Figure S3**: Self-assembly mechanisms of hybrid tumoroids. **A)** Quantification of DSLB-based synthetic cells integration into tumoroids formed from Panc-1 cells. DSLBs were either incubated with singularized synthetic cells for 48 hours (co-incubation), or added to pre-formed spheroids (sequential incubation). Results shown as mean $\pm$SD from n>23, *** $p<0.001$ unpaired two-tailed t-test. **B)** SDS PAGE of serum opsonization levels from the synthetic cell panel incubated in 10% fetal bovine serum for 1 hour. Total band intensity for each lane is indicated in the bottom. **C)** Quantification of DSLB-based synthetic cells integration into tumoroids formed from Panc-1 cells either with plain DSLBs or DSLB produced with 10 mol% PEG5000 on the lipid bilayer surface. Results shown as mean $\pm$SD from n>20, p values as indicated from unpaired two-tailed t-test. **D)** Quantitative PCR of Bfl-1 RNA levels in Panc-1 hybrid tumoroids formed with DSLBs presenting CD40L, plain DSLBs or no DSLBs. Results are shown from 3 separate replicates shown as mean $\pm$SD, normalized to RNA levels of glyceraldeyde-3-phosphate dehydrogenase. P values as indicated from one-way ANOVA.

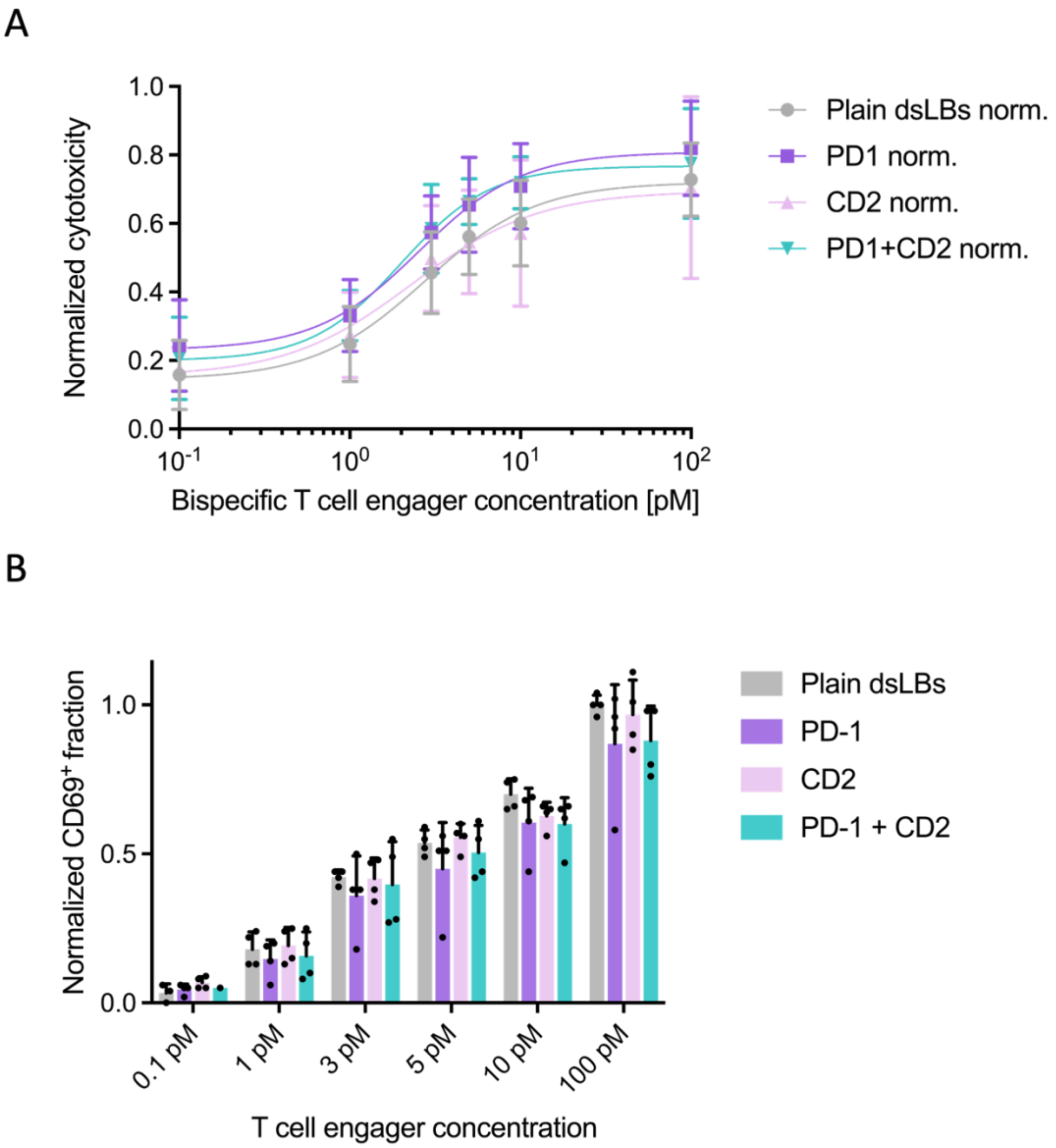


**Figure S4** Bispecific T cell engager-mediated killing of hybrid tumoroids. **A)** Lactate dehydrogenase release assay for quantification of killing of ART-TIME containing Panc-1 hybrid spheroids by primary human CD8 T cells in a bispecific T cell engager dilution series. T cells were incubated with hybrid tumoroids preformed for 48 hours at a 5:1 T cell to cancer cell ratio. Results are shown from n=2 donors as mean ±SD from two replicates each. **B)** Flow cytometry quantification of the fraction of CD69 expressing primary human CD8 T cells incubated with ART-TIME containing Panc-1 3D cultures for 12 hours in the presence of increasing bispecific T cell engager concentrations. Results are shown from n=2 donors as mean ±SD from two replicates each.

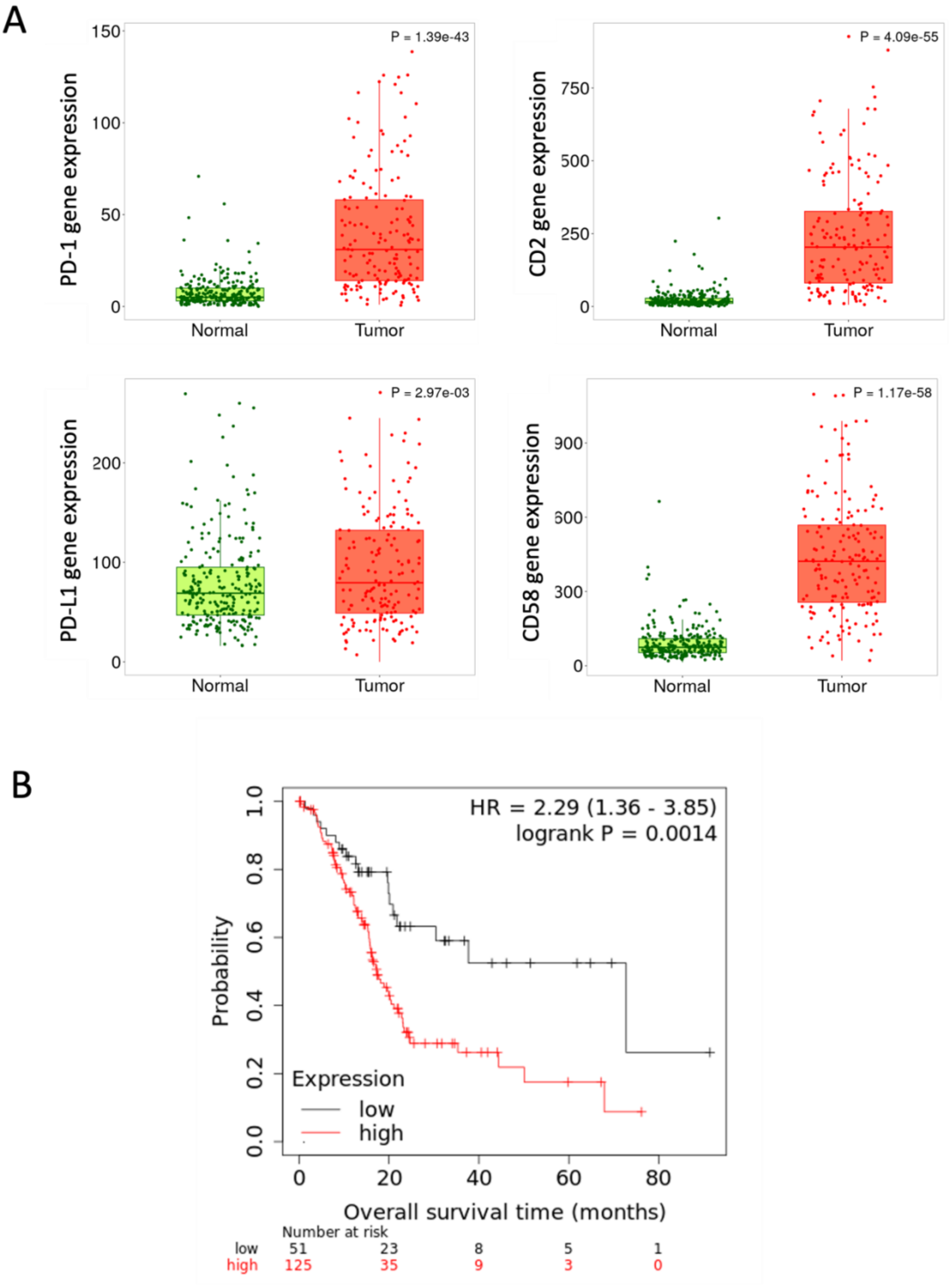


**Figure S5** Clinical relevance of PD-1/PD-L1 and CD2/CD58 expression. RNA expression levels of PD-1 and PD-L1 as well as CD58 and CD2 from 177 patients samples quantified from tumorous tissue and 252 normal pancreatic tissues. **B)** Survival analysis in pancreatic adenocarcinoma samples using the combined expression of PDL1 and CD58 and overall survival data shows markedly worse outcome in those with higher expression.
